# The emergence of human gastrulation upon *in vitro* attachment

**DOI:** 10.1101/2023.05.16.541017

**Authors:** Riccardo De Santis, Eleni Rice, Gist Croft, Min Yang, Edwin A. Rosado-Olivieri, Ali H. Brivanlou

## Abstract

While studied extensively in model systems, human gastrulation remains obscure. This process starts upon blastocyst implantation into the uterine wall, which is assumed to occur after 14 days post-fertilization. The scarcity and limited access to fetal biological material as well as ethical considerations limit our understanding of the cellular and molecular portrait of human gastrulation. *In vitro* culture of natural human blastocysts shed light on the second week of human development, unveiling an unexpected level of self-organization embedded in the pre-gastrulating embryo, yet whether they can undergo gastrulation upon *in vitro* attachment remains elusive. Blastocyst models called *blastoids*, which are derived from human pluripotent stem cells, provide the opportunity to reconstitute post-implantation human development *in vitro* with unlimited biological material. Here we show that human *blastoids* break symmetry and undergo gastrulation upon *in vitro* attachment. scRNA-seq of these models replicate the transcriptomic signature of the natural human gastrula, recapitulating aspects of the second to the third week of human development. Surprisingly, analysis of developmental timing reveals that in both *blastoid* models and natural *in vitro* attached human embryos, the onset of gastrulation as defined by molecular makers, can be traced to time scales equivalent to 12 days post-fertilization, which appeals for a reconsideration of the onset of human gastrulation upon extended *in vitro* culture.

## Introduction

*Gastrulation* is one of the most dramatic moments in our early life: a time when in the implanted embryo, a sheet of cells called *epiblast* that gives rise to the entire embryo, breaks symmetry, undergoes morphogenetic movements, and establishes the primordia of the three-germ layers and body axis. This process can be recognized by the emergence of morphological landmarks such as the formation of the organizer in Teleost and Amphibians, or node and primitive streak in birds and mammals (Solnica-Krezel and Sepich, 2012; Ghimire et al., 2021). However, the molecular program initiating gastrulation occurs before the emergence of the morphological landmarks as demarcated by the expression of BRACHYURY (BRA)(Rivera-Pérez and Magnuson, 2005).

The establishment of culture systems that allowed *in vitro* attachment of natural human blastocysts (i.e., blastocysts obtained from fertilization) shed light on the second week of human development and unveiled an unexpected level of self-organization embedded in the pre-gastrulating embryo (Deglincerti et al., 2016; Shabazi et al., 2016). These embryos, however, did not display the morphological signatures of gastrulation. We therefore asked whether the onset of human gastrulation can be recognized by the expression of molecular markers before the emergence of morphological features. It is self-evident that, regardless of rules and guidelines, the sheer number of human embryos required to achieve the same level of scrutiny as in model systems is unrealistic. To alleviate this inevitable limitation, *in vitro* models of the human gastrula derived from human pluripotent stem cells (hPSCs) were recently developed (Warmflash et al., 2014; Shao et al., 2017; Martyn et al., 2018; Simunovic et al., 2019; Zheng et al., 2019; Moris et al., 2020, Simunovic et al., 2022; reviewed in Shahbazi, 2020; van den Brink and van Oudenaarden, 2021). However, these systems do not self-organize from a blastocyst-like structure, lack preimplantation extraembryonic cells, and rely on the exogenous presentation of morphogens such as BMP4, WNT, and Activin A.

Recently, naïve hESCs were shown to be able to self-organize into 3D blastula models (*blastoids)* that attach *in vitro* and recapitulate features of human post-implantation development until 13 dpf, in the absence of cellular and morphological manifestations of primitive streak formation (Yu et al., 2021; Yanagida et al., 2021; Kagawa et al., 2022). Following these pioneering studies, we show that human *blastoids* generated from naïve hESCs can gastrulate and self-organize to generate embryonic and extraembryonic germ layers upon *in vitro* attachment, reconstituting the molecular signature of the human primitive streak. Our study traces the origin of human gastrulation to 12 dpf in *in vitro* attached natural human embryos, and provides the *in vitro*-reconstituted transcriptomic signature of early human development.

## Results

### *in vitro* attached human *blastoids* gastrulate

RUES2 (NIHhESC-09-0013) cells were reprogrammed from post-implantation “primed” to a pre-implantation “naïve” state by chemical resetting (Guo et al., 2017). This was manifested by a change in colony morphology (from flat epithelial to dome-shaped) and expression of preimplantation epiblast markers KLF17 and SUDS2 (Suppl. Fig 1A-B) (Bredenkamp et al., 2019). Upon differentiation in microwells (Aggrewells) using PALLY-LY media (Kagawa et al 2022), naïve RUES2 cells self-organized into 3D *blastoids* (Fig 1A, Suppl. Fig 1C), harboring epiblast (OCT4^+^), primitive endoderm (SOX17^+^), and trophectoderm (GATA3^+^) populations (Fig 1B) (Kagawa et al., 2022). We modified our *in vitro* attachment protocol for natural human embryos (Deglincerti et al., 2016; Shabazi et al., 2016) to include conditions developed for the maintenance of extended culture of human and non-human primate embryos (Ma et al., 2019; Niu et al., 2019; Xiang et al., 2019). RUES2 *blastoids* were attached on plastic substrates coated with Laminin-521, in the presence of ROCK-inhibitor (Y-27632) and extracellular matrix in the media (geltrex 5%) (Material and Methods). Epiblast symmetry breaking and initiation of primitive streak and mesoderm formation were evaluated by expression of OCT4^+^ (epiblast/ectoderm), GATA6^+^ (endoderm) and OCT4^+^/BRA^+^ (primitive streak) after 7 days post-attachment (dpa). We use dpa instead of dpf because hPSCs do not have a time zero as identified by fertilization, upon which time is counted in natural human embryos. At dpa 7, a subset of *blastoids* displayed a small clump of OCT4^+^/BRA^+^ cells, matching the molecular signature of an early primitive streak (Fig 1C). Immunostaining analysis of *in vitro* attached *blastoids* at 10 dpa resulted in a prominent BRA^+^ territory in a fraction of *in vitro* attached *blastoids* (Fig 1D). Both OCT4^+^ and OCT4^+^/BRA^+^ populations increased over time, suggesting an enlargement of the epiblast population, as well as induction and expansion of primitive streak and mesoderm populations (Fig 1E, N=51 from 3 independent biological replicates). The fraction of *blastoids* that broke epiblast symmetry expanded over time and was 8% at 7 dpa and 52% at 10 dpa (Fig 1F). A BRA^+^ population was induced across a wide range of OCT4^+^ epiblast areas, suggesting that the absolute size of the epiblast was not associated with epiblast symmetry breaking (Fig 1G; Suppl. Fig 2A). Self-organization of a gastrulating epiblast and symmetry breaking was also observed in *blastoids* derived from another human naïve pluripotent stem cell line, niPSC75.2 (Suppl. Fig 2B). We conclude that *in vitro* attached human *blastoids* self-organize an OCT4^+^/BRA^+^ human primitive streak population in 7-10 dpa.

**Fig 1:**
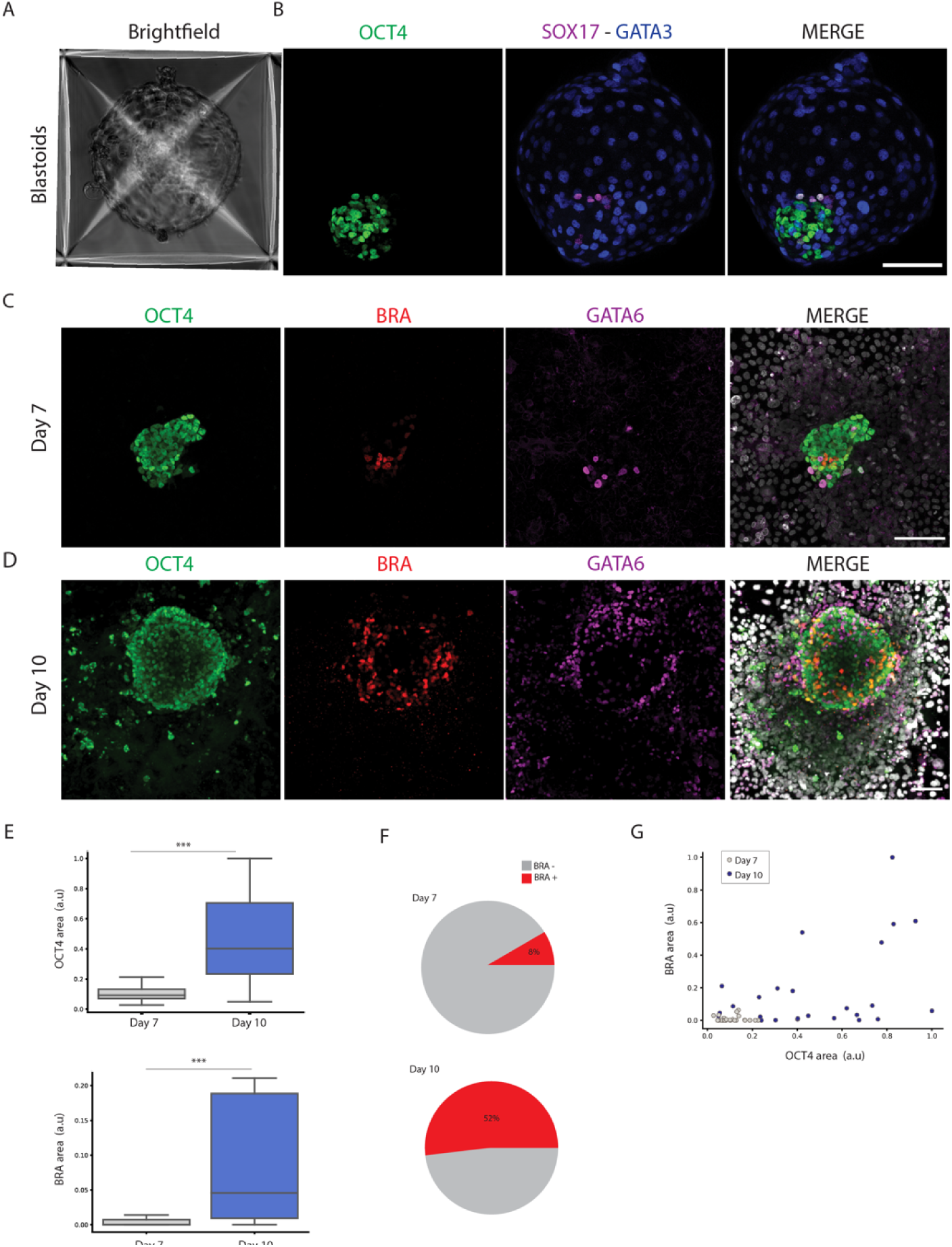
*In vitro* attached human *blastoids* break epiblast symmetry and self-organize primitive streak and mesoderm populations. A) Brightfield image of PALLY-LY differentiated *blastoids* in Aggrewells. B) Immunostaining *blastoids* displays OCT4 (green), SOX17 (magenta), and GATA3 (blue) marker genes distinguishing epiblast, primitive endoderm, and trophectoderm. Scale bar 100um. C) *In vitro* attached human *blastoids* analyzed by immunostaining show OCT4^+^/BRA^+^ cells 7 dpa. OCT4^+^ cells label the epiblast territory, GATA6^+^ the primitive endoderm, and BRA^+^ the primitive streak and mesoderm populations. Immunostaining pseudo colors: OCT4 (green), BRA (red), GATA6 (magenta), and merge with DAPI (grey). Scale bar: 100um. D) Immunostaining at 10 dpa displays OCT4^+^-epiblast, BRA^+^-primitive streak and mesoderm, and GATA6^+^-endoderm cells. Immunostaining pseudo colors: OCT4 (green), BRA (red), GATA6 (magenta), and merge with DAPI (grey). Scale bar: 100um. E) Scaled area quantification of OCT4^+^ and BRA^+^ cells at 7 and 10 dpa. Data are displayed as a boxplot (center line displays the median, and box limits are the 25th and 75th percentiles) (n = 51 from 3 independent differentiations). Student’s t-test, unpaired with two tails. ***p<0.001. F) Pie chart of *in vitro* attached *blastoids* that broke symmetry at 7 and 10 dpa. G) Area quantification is displayed as a scatterplot. x-axis reports the scaled OCT4 while the y-axis is the BRA scaled area. Each dot represents an individual attached *blastoid* that has been color-coded according to the time of analysis. Grey dots represent *blastoids* 7 dpa and blue dots *blastoids* at 10 dpa.

### scRNA-seq of *in vitro* attached human *blastoids* reveal convergent features with human gastrula

Two different time points (7 and 10 dpa) were analyzed by scRNA-seq. 16 distinct clusters comprised of derivatives of the three germ layers and extraembryonic cells were identified (Fig 2 A-B, Suppl. Fig 3A-B). Cell type annotation was based on the expression of markers genes and revealed the presence of epiblast, ectoderm, primordial germ cells (PGCs), two distinct populations of the primitive streak, three mesodermal sub-populations, cardiac mesoderm, endothelial progenitors, definitive endoderm, hypoblast, yolk sack endoderm, cytotrophoblast, syncytiotrophoblast, and amniotic cells (Fig 2C, Suppl. Fig 3C, Table 1). Immunostaining analysis confirmed the presence of MIXL1^+^, GATA4^+^, HAND1^+^, FOXA2^+^, and SOX17^+^ cells, distinguishing primitive streak, mesoderm, and endoderm populations in *in vitro* attached blastoids at 10 dpa (Fig 2D). In addition to derivatives of three germ layers, immunostaining analysis displayed trophoblast cells (GATA3^+^/KRT7^+^) and an amnion (ISL1^+^) population around the epiblast territory, although no prominent amniotic cavity was detected at 10 dpa (Suppl. Fig 4A-B). Furthermore, scRNA-seq analysis distinguished the molecular signature of anterior and posterior primitive streak populations, with the anterior population resembling the Spemann organizer (GSC^+^/CER1^+^/FST^+^/NOG^+^) (Suppl. Fig 5A-B). Thus, single cell transcriptomic of *in vitro* attached human *blastoids* revealed the presence of derivatives of the three germ layers and extraembryonic cellular populations at 7-10 dpa.

**Fig 2:**
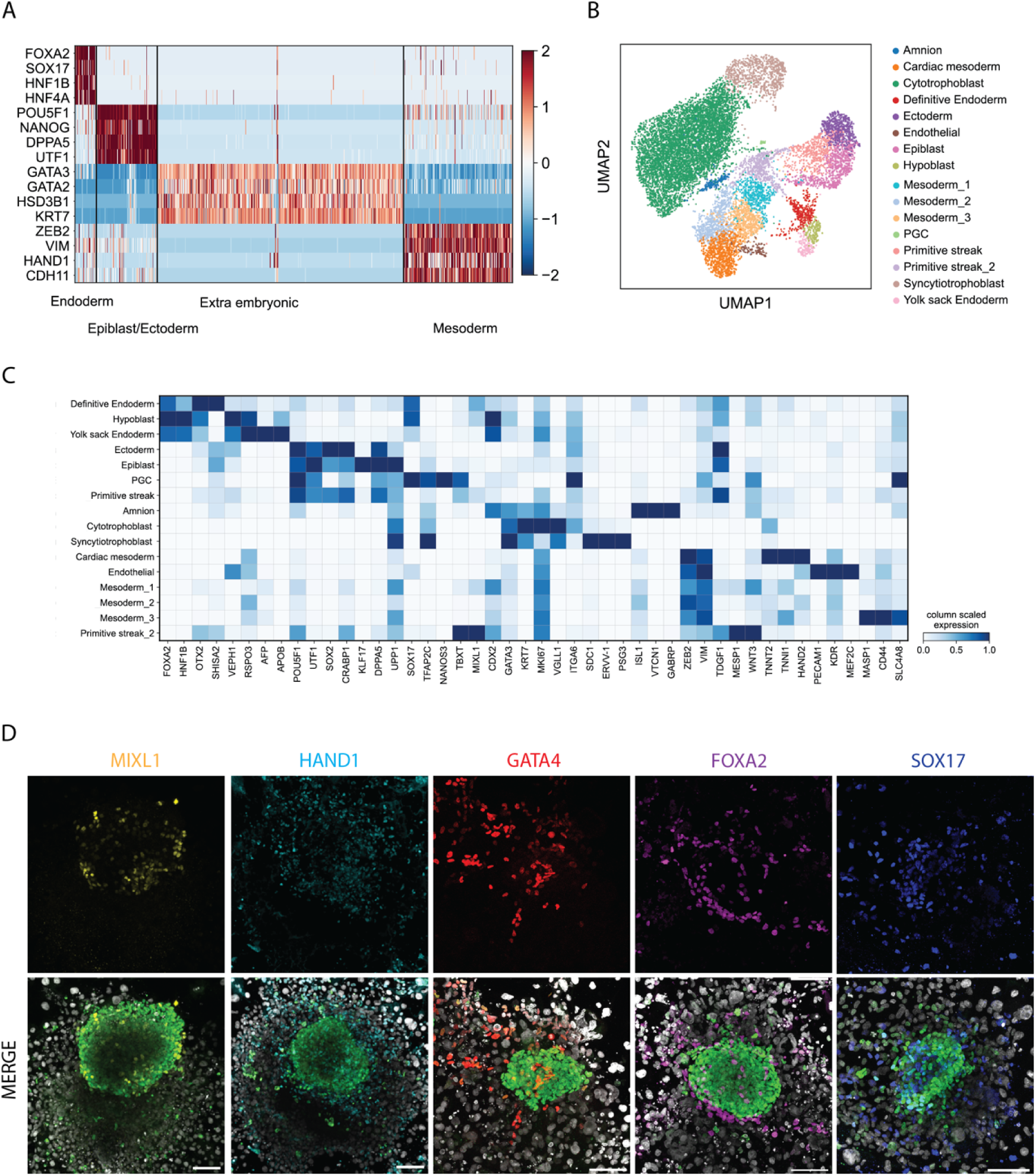
Single cell transcriptomics of *in vitro* attached human *blastoids* distinguished embryonic and extraembryonic populations in presence of self-organized signaling dynamics. A) Heatmap displays z-score scaled expression values for marker genes that distinguish extraembryonic (GATA3, GATA2, HSD3B1, KRT7), epiblast/ectoderm (POU5F1, NANOG, DPP5, UTF1), mesoderm (ZEB2, VIM, HAND1, CDH11), and endoderm (FOXA2, SOX17, HNF1B, HNF4A) populations at 7 and 10 dpa (7 dpa: 7354 cells and 10 dpa: 5489 cells). B) UMAP plot shows the identified cell populations: cytotrophoblast, syncytiotrophoblast, amnion, epiblast, ectoderm, primordial germ cells, primitive streak 1-2, mesoderm 1-2-3, cardiac mesoderm, endothelial, definitive endoderm, hypoblast, and yolk sack endoderm. C) Heatmap of average scaled expression displaying marker genes that distinguish individual cell populations. D) Immunostaining of 10 dpa human *blastoids.* OCT4 (green) and DAPI (grey) are used to identify *blastoids*. MIXL1 (yellow), HAND1 (cyan), GATA4 (red), FOXA2 (magenta), and SOX17 (blue) identify mesoderm and endoderm populations. Scale bar: 100um.

We then compared our dataset to the recently published scRNA-seq transcriptome profile of the *in vivo* gastrulating human embryo during the third week of development (16-19 dpf) (Tyser et al., 2021). Datasets integration displayed preferential alignment of the human gastrula within the epiblast/ectoderm, mesoderm, and endoderm clusters (Fig 3A). Cluster level correlation analysis showed several *blastoid* cell types displaying high correlation with distinct *in vivo* populations (Fig 3B). Hierarchical clustering identified multiple subgroups containing *in vitro* cell populations and their corresponding *in vivo* counterparts (Fig 3C), suggesting cell type-specific convergent gene expression between *in vitro* attached *blastoids* and *in vivo* gastrulating human embryos. To further corroborate our classification, we harnessed an embryonic cell classifier to validate our dataset and cell type classification (Zhao et al., 2021). The classifier predicted cell types that match our marker-based cell type classification (Suppl. Fig 6A, Table 2). To further support our analysis, we integrated our scRNA-seq with published *in vivo* mouse and monkey datasets (Pijuan-Sala et al., 2019; Zhai et al., 2022), and with *in vitro* cultured human and monkey embryos (Ma et al., 2019; Xiang et al., 2019),), that altogether displayed convergent gene expression matching cellular populations in embryonic and extraembryonic tissues of *in vitro* attached human *blastoids* (Suppl. Fig 7A-B; Suppl. Fig 8A-B). In conclusion, single cell transcriptomics of *in vitro* attached human *blastoids* displayed the molecular signatures of embryonic and extraembryonic populations that match *in vivo* populations, including those of the human gastrula in the third week of development.

**Fig 3:**
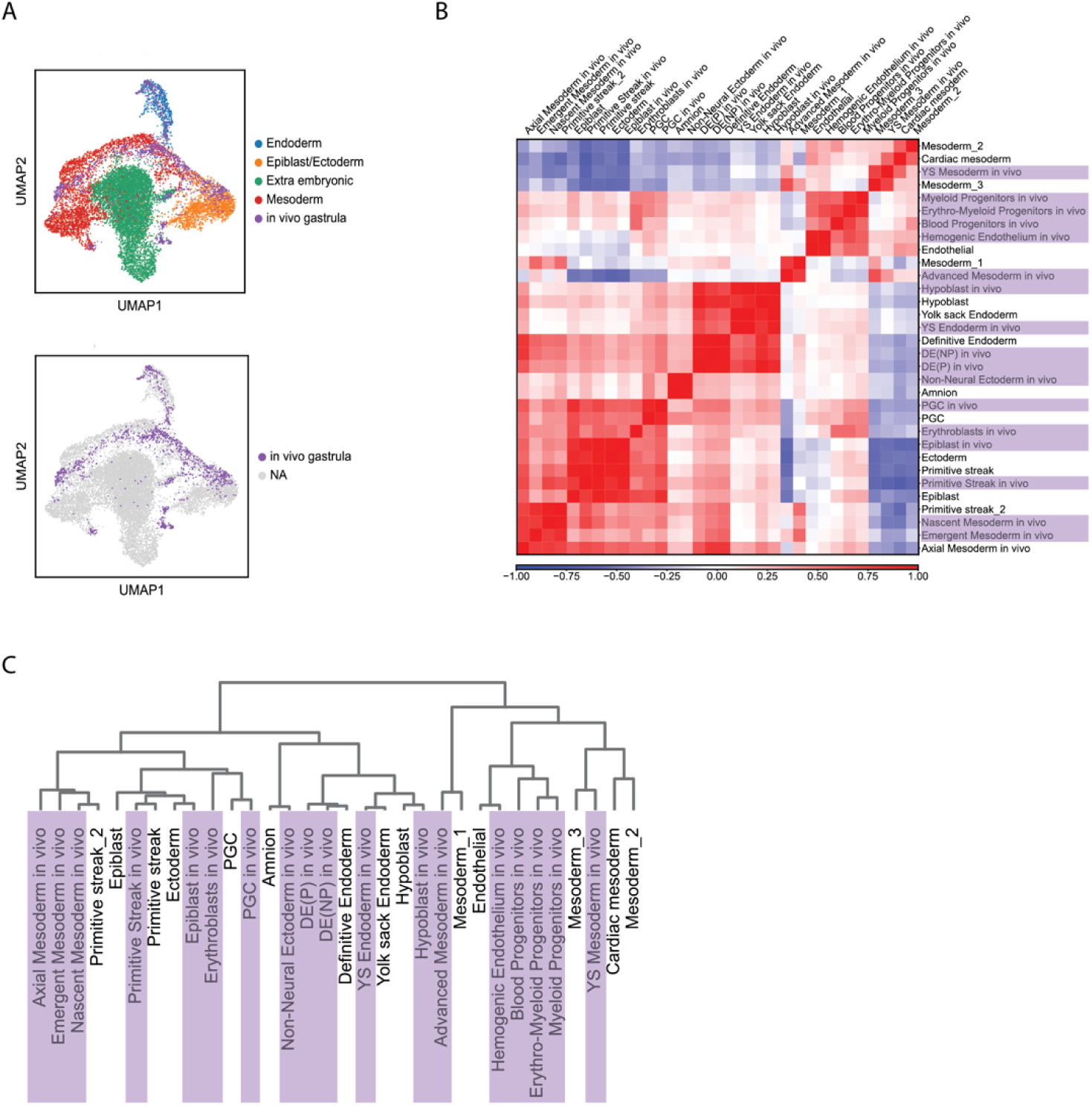
In-vitro attached human *blastoids* recapitulate the cellular populations of the human gastrula. A) UMAP shows the integration of 7-10 dpa *blastoids* scRNA-seq and the human gastrula dataset at 16-19 dpf (Tyser et al., 2021). *In vitro* attached *blastoids* are color coded according to their lineage. The human gastrula dataset preferentially integrates with the epiblast/ectoderm, mesoderm, and endoderm lineages. B) Correlation analysis using the commonly detected highly variable genes between the human gastrula dataset (Tyser et al., 2021) and the *in vitro* attached human *blastoids* dataset displayed as a z-score heatmap. The clusters from the human gastrula dataset are shown with their original annotations followed by an “*in vivo”* label and are highlighted in light purple. C) Hierarchical clustering is displayed as a dendrogram. The human gastrula (Tyser et al., 2021) labels are highlighted in light purple.

The onset of gastrulation is regulated by the signaling cascade that starts with BMP4, which then induces WNT that in combination with NODAL controls mesoderm and endoderm differentiation (Arnold and Robertson, 2009; Warmflash et al., 2014; Morgani et al., 2018; Martyn et al., 2018). scRNA-seq analysis of *in vitro* attached human *blastoids* displayed endogenous expression of WNT3 and BMP4 in the amnion and in the mesodermal clusters, while NODAL expression was mostly confined to the epiblast, ectoderm, endoderm and primitive streak populations (Suppl. Fig 9A). In agreement with cell type-specific ligand expression, we also observed induction of ID2 and LEF1 gene expression, which are immediate-early target genes of the BMP and WNT pathways, suggesting self-organized active signaling (Suppl. Fig 9B)(Sasaki et al., 2016; Martyn et al., 2019). Treatment with small molecules inhibiting BMP (LDN-193189) or WNT (IWR1-endo) pathways starting from 5 dpa, significantly reduced BRA expression demonstrating BMP- and WNT-dependent induction of primitive streak and mesoderm populations upon *in vitro* attachment (Suppl. Fig 9C). Collectively, these data show that *in vitro* attached *blastoids* self-organize BMP, WNT and NODAL signaling pathways in a cell type-specific manner, controlling the emergence of primitive streak and mesoderm populations upon *in vitro* attachment.

### *in vitro* attached natural human embryos break epiblast symmetry at 12 dpf

*In vitro* attached human *blastoids* displayed epiblast symmetry-breaking, marked by BRA expression in the epiblast territory, around 7 dpa (Fig 1C). We matched this early sign of epiblast symmetry-breaking with the timeline of *in vitro* attached natural human embryos for analysis at 12-13 dpf. As with the *blastoids*, *in vitro* attachment was performed on functionalized plastic surfaces with laminin 521 and immunostaining analysis of cell type-specific markers was performed at 12 dpf. OCT4 staining was used to identify the epiblast population, GATA6 for primitive endoderm, BRA for primitive streak and mesoderm, and F-Actin and DAPI to mark the embryo structure (Fig 4A). Among the OCT4^+^ epiblast cells, we revealed a subset of cells that were also positive for BRA and negative for GATA6, indicative of an early primitive streak and mesoderm population at 12 dpf (Fig 4B). 3D rendering of the *in vitro* attached human embryo localized the OCT4^+^/BRA^+^ population on one side of the epiblast, suggesting the establishment of an anterior-posterior axis in a natural human embryo at 12 dpf (Fig 4C). We further investigated whether the use of laminin 521 functionalized plastic substrates for *in vitro* attachment was required for epiblast symmetry breaking. Human embryos were *in vitro* attached according to Deglincerti et al., in the absence of functionalized substrates and consistently displayed self-organization of a BRA^+^ population at 12 dpf (Suppl. Fig 10).

**Fig 4:**
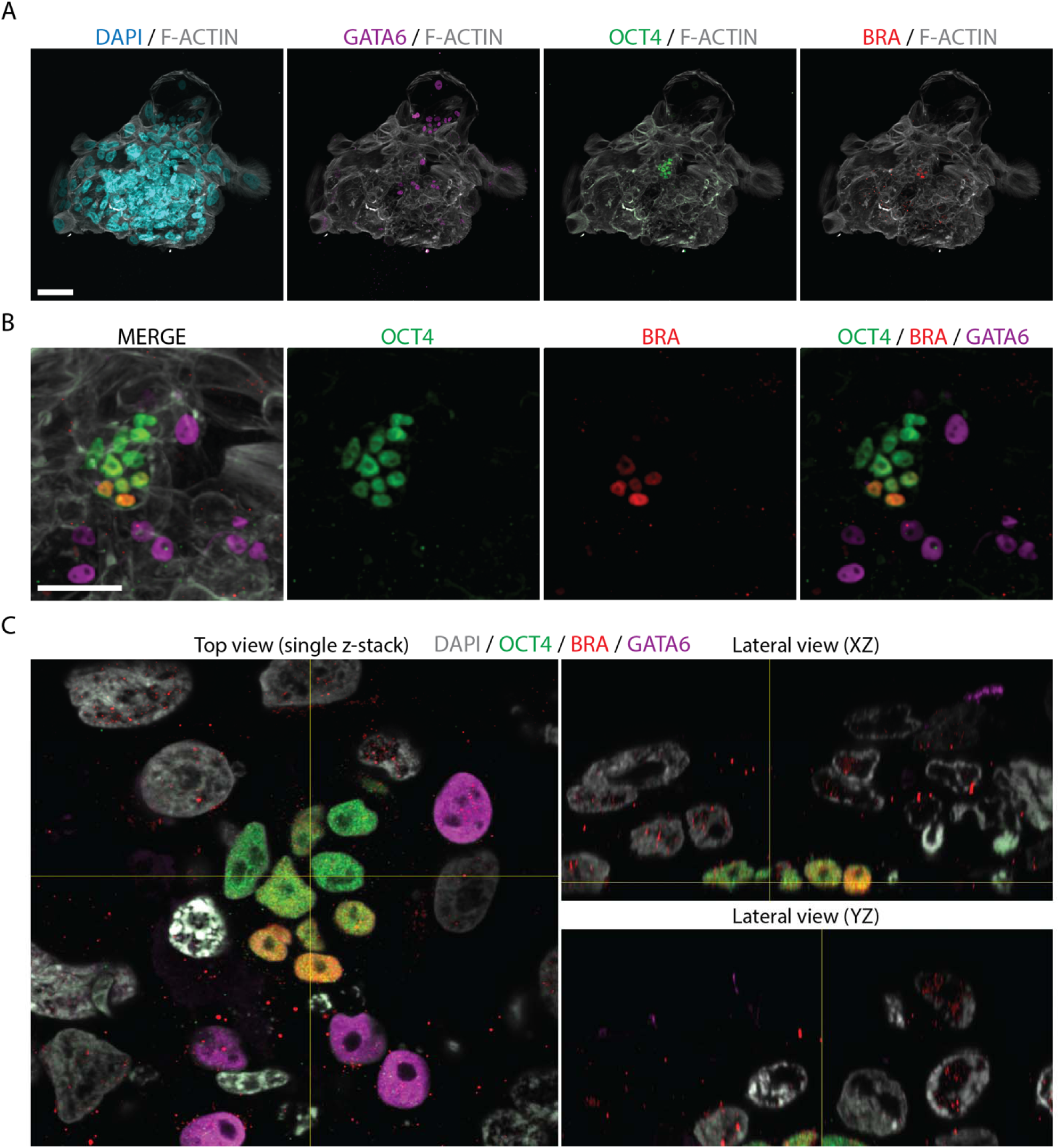
*In vitro* attached human embryos break epiblast symmetry and self-organize a BRA-positive axial population at 12 dpf. A) Immunostaining of the *in vitro* attached human embryo at 12 dpf. A low-magnification field of view shows DAPI (cyan), GATA6 (magenta), OCT4 (green) and BRA (red) signals, all overlayed on the F-ACTIN (grey) staining. OCT4 localizes the epiblast territory and GATA6 the primitive endoderm cells. BRA is used as a primitive streak and mesoderm marker. Scale bar: 100um. B) Immunostaining of the *in vitro* attached human embryo at 12 dpf. Zoomed-in field of view shows the merged image of OCT4 (green), BRA (red), GATA6 (magenta) and F-ACTIN (grey) signals. Scale bar: 50um. C) 3D orthogonal projections of the *in vitro* attached human embryo at 12 dpf. Top and lateral views show the merge of OCT4 (green), BRA (red), GATA6 (magenta), and DAPI (grey). Scale bar: 50um.

In all, *in vitro* attached human embryos displayed the earliest sign of primitive streak and mesoderm differentiation at 12 dpf, breaking epiblast-symmetry and self-organizing an anterior-posterior axis marked by expression of BRA in the epiblast territory.

## Discussion

We have shown that upon *in vitro* attachment, natural human embryos and synthetic models of human blastocysts (*blastoids)* progress through stages of post-implantation development, including the onset of primitive streak and mesoderm formation, modeling key aspects of human gastrulation *in vitro*. Taking advantage of stem cell-derived human *blastoids,* we defined the timeline of epiblast symmetry breaking upon *in vitro* attachment, discovering the earliest molecular sign of primitive streak formation at 12 dpf *in vitro* attached human embryos. Detection of primitive streak molecular markers before 14 dpf, suggests an earlier time for the onset of gastrulation upon extended culture *in vitro*. It is tempting to speculate that this event also occurs during human development *in vivo*, but without the precise staging of *in vivo* human samples, which is ethically and practically impossible, it is not possible to confirm directly. On the other hand, *in vitro* culture might impact the timeline of epiblast symmetry breaking. Our data invites for reconsideration of current guidelines and regulations, suggesting temporal staging of *in vitro* cultured human embryos and embryoids using molecular markers instead of morphological criteria or a priori-defined timelines. This evidence is also supported by recent data obtained from *in vitro* cultured non-human primate embryos, that displayed BRA expression and the onset of a primitive streak population around 11-13 dpf upon *in vitro* attachment (Niu et al., 2019; Ma et al., 2019).

Extended culture of *in vitro* attached human *blastoids* up to 10 dpa resulted in the self-organization of derivatives of the three germ layers, in the absence of exogenous BMPs and WNTs ligands or cell mixing with other extra-embryonic cell types. Single-cell transcriptomics of self-organizing *in vitro* attached human *blastoids* allowed temporal and cellular matching with the dataset of the *in vivo* human gastrula at day 16-19 dpf (Tyser et al., 2021). *In vitro* attached *blastoids* recapitulated most of the cellular populations present in the human gastrula; however, while mesodermal populations displayed proximal-distal organization from the epiblast territory, they did not self-organized the morphological features of an *in vivo* primitive streak at this stage. Attachment on 2D plastic substrates may represent one of the limiting factors for further embryonic development. We envision that the next generation of models will rely on biocompatible polymers capable of mimicking implantation while also supporting expansion in 3D, allowing self-organization and *in vivo*-like morphogenesis of differentiated tissues and organs. It has been recently shown that taking advantage of hydrogel embedding and optimized culture conditions, 3D monkey embryos develop *in vitro* up to 25 dpf, recapitulating the cellular and molecular features of Carnegie stage 8 monkey embryos, including early organogenesis (Gong et al., 2023), Interestingly, it has also been recently shown that assembled 3D mouse *embryoids,* derived from mouse naïve pluripotent stem cells, are able to progress through stages of post-implantation development, including organogenesis, in the absence of physical attachment on a substrate (Tarazi et al., 2022; Amedei et al., 2022). Similarly, monkey blastocysts cultured under optimized 3D culture conditions have been shown to progress *in vitro* up to 25 dpf, self-organizing derivatives of the three germ layers and forming neural tube-like structures with antero-posterior and dorso-ventral embryonic axes (Zhai et al., 2023). These findings suggest that physical attachment on a substrate may not be required for further progression *in vitro*, whereby extraembryonic tissues and/or optimized 3D culture conditions are in place to sustain embryos and *embryoids* self-organization.

Ascertaining the cellular and molecular logic of human gastrulation has tremendous implications for understanding the principles of body axes formation and potential causes of early miscarriages linked to embryonic failure. It has been estimated that up to 30% of total miscarriages occur after the second week of gestation (Ghimire et al., 2021), coincidentally aligning with the onset of gastrulation and possibly linked to problems in establishing axial organization during embryonic development. Our current knowledge of the mechanisms directing axis formation in the context of humans is limited and derives mostly from animal models. Important parameters such as embryo geometry, time of gestation, and gene regulatory networks are species-specific attributes that must be considered when mechanisms of development are extrapolated from animal models. Studies on *in vitro* cultured natural human embryos and fetal samples provide an experimental framework to evaluate and interpret the cellular and molecular players involved in human gastrulation. Complementary to natural human embryos and fetal samples, models based on human stem cells offer virtually unlimited biological material, extensive possibilities for genetic manipulation, and experimental investigation. Although we trust that stem cell-derived synthetic models will become more representative of *in vivo* situations, we also believe that comparisons with actual fetal samples, *in vitro* cultured natural embryos, and cross-species evaluations represent essential steps to evaluate *in vitro* stem cell-based models.

## Author contributions

RDS conceived the project, performed human embryos and *blastoids* experiments, analyzed the data and wrote the manuscript. ER contributed to human pluripotent stem cell culture, *blastoids* generation and immunostaining. GC performed human embryo experiments. MY recruited human embryos and contributed to human embryo experiments. EARO contributed to scRNA-seq analysis and revised the manuscript. AHB conceived and coordinated the project, wrote the manuscript, and secured funding.

## Highlights

*-In vitro* attached human *blastoids* break epiblast symmetry and self-organize primitive streak and mesoderm populations

-Single cell transcriptomic of *in vitro* attached human *blastoids* recapitulate the cellular populations of the human gastrula

-*In vitro* attached natural human embryos break epiblast symmetry and self-organize a BRA-positive axial population at 12 dpf.

## Acknowledgments

We are grateful to Brivanlou lab members for their input and criticism. We would like to thank the Rockefeller Bio-Imaging Resource Center for imaging technical support and the NYU Genomic Core Facility for sequencing technical support. We also would like to thank Dr. Norbert Gleicher at the Center for Human Reproduction and Dr. Pasquale Patrizio at Yale Fertility Center for providing donated human embryos from patients for this study, and Dr. Emanuela Molinari for helping with embryo handling. R.D.S. was supported by EMBO-LTF-254-201.

## Ethics statement

The research described in this manuscript was approved by The Rockefeller University Institutional Review Board according to university policy for research involving human embryonic stem cells and human embryos, and governed by a Tri-Institutional Stem Cell Initiative, Embryonic Stem Cell Oversight (Tri-SCI ESCRO) Committee that oversees human Embryo Research Oversight (EMRO) and serves The Rockefeller University, Memorial Sloan Kettering Cancer Center, and Weill Cornell Medicine. The ESCRO committee is an independent group charged with oversight of research involving human pluripotent stem cells and embryos to ensure compliance with university policies. The Tri-SCI ESCRO committee is composed of members with scientific and bioethical expertise. Established ESCRO protocols are reviewed annually and adhere to ISSCR 2021 guidelines. Supernumerary IVF-derived human embryos donated for research were obtained from the Center for Human Reproduction and the Yale Fertility Center. Embryo donation was facilitated through an informed consent process according to NAS Guidelines for Human Embryonic Stem Cell Research and the Tri-Institutional Stem Cell Research Operating Procedures for ESCRO Reviewed Research.

## Material and methods

Human pluripotent stem cell culture and chemical reprogramming RUES2 hESCs (NIHhESC-09-0013) were maintained in primed state using MEF-conditioned media (CM) supplemented with FGF2 (20ng/mL) on Geltrex coated dishes (ThermoFisher Scientific). CM-FGF2 media was refreshed every day, cells were passaged every 3-5 days using Accutase or ReleSR (ThermoFisher Scientific).

Reprogramming to a naïve state was performed according to Guo et al., 2017, Bredenkamp et al., 2019 with minor modifications. In brief, undifferentiated prime RUES2 cells (in CM-FGF2) were dissociated as single cells with Accutase (Stem Cell Tecnologies), and 100,000 cells were plated on MEF plates with the addition of ROCK-inhibitor (10uM). The following day, CM-FGF2 media was refreshed allowing cells to form colonies. Two days after seeding, reprogramming was started using N2B27 media (50% DMEM-F12, 50% Neurobasal, GlutaMAX 1X, beta-mercaptoethanol 55mM, N2 0.5X and B27 0.5X), supplemented with human LIF (10 ng/mL), PD0325901 (1uM) and VPA (1uM) (all from Stem cell technologies). 50% of the media was refreshed two days after, and on day three media was switched to PXGL media (N2B27 media supplemented with PD0325901 1uM, XAV939 2uM, Go6983 2uM and LIF 10ng/mL) and refreshed every day until day 10.

Human naïve pluripotent stem cells were maintained up to 25 passages in PXGL media. Naïve cells were passaged on MEF plates upon single cell dissociation with Accutase every 3-5 days in PXGL media supplemented with ROCK-inhibitor (10uM) and Geltrex (1%).

### *Blastoids* differentiation in microwells

Naïve human pluripotent stem cells were differentiated to form *blastoids* according to Kagawa et al., 2022a; Kagawa et al., 2022b with minor adjustments. In brief, naïve pluripotent stem cells were dissociated as single cells using Accutase, and MEF were removed upon differential attachment on gelatin plates for 45 minutes. Unattached human naïve cells were collected, counted, and seeded overnight on Aggrewells 400 at the density of 160 cells/well in N2B27 media supplemented with ROCK-inhibitor (10uM). The day after, media was switched to N2B27 media supplemented with PALLY (PD0325901 1uM, A83 1uM, LPA 0.5uM, LIF 10ng/mL, and ROCK-inhibitor 10uM) for 48 hours. Following 48 hours of PALLY media, media was refreshed for an additional 48 hours with N2B27 media supplemented with LY (LPA 0.5uM and ROCK-inhibitor 10uM). Based on morphological criteria, *blastoids* were picked manually for further experiments.

### Human *blastoids in vitro* attachment protocol

*Blastoids* were selected based on morphological criteria and transferred on IBIDI 8 wells coated with Laminin-521 (10ug/mL) (Biolamina) in IVC1 media adapted from Deglincerti et al., 2016, and Xiang et al., 2019. IVC1 media was composed of 50% DMEM/F12 (Thermo Fisher Scientific), 50% Neurobasal (Thermo Fisher Scientific) supplemented with 20% (vol/vol) heat-inactivated fetal bovine serum (FBS, R&D S1115050H), GlutaMAX (1X; Thermo Fisher Scientific), 1× ITS-X (Thermo Fisher Scientific), 8 nM β-estradiol, 200 ng/ml progesterone, and 25 μM N-acetyl-L-cysteine (all from Sigma Aldrich). Two days post attachment, media was switched to IVC2 media: 50% DMEM/F12 (Thermo Fisher Scientific), 50% Neurobasal (Thermo Fisher Scientific) supplemented with 30% KnockOut Serum Replacement (10828028, Thermo Fisher Scientific), GlutaMAX (1X; Thermo Fisher Scientific), 8 nM β-estradiol, 200 ng/ml progesterone and ROCK-inhibitor (10uM). IVC2 media was refresh every other day. Extended culture up to day 10 was obtained upon extracellular matrix supplementation in IVC2 media starting from 5 dpa with 5% Geltrex (Thermo Fisher Scientific).

### Human embryo thawing and *in vitro* attachment protocol

Cryopreserved IVF-derived human embryos (5-6 dpf) were thawed using the Kitazato kit (VT602US, Kitazato USA) according to manufacturer instructions. In brief, straws containing individual embryo were placed into TS1 solution for 1 minute, passed twice in DS2 solution for 2 minutes and twice in WS3 solution for 3 minutes. Embryos were cultured overnight in GLHP media drops (GlobalHP) under mineral oil (FUJIFILM). The day after, hatched blastocysts were transferred on Laminin-521 coated IBIDI 8 wells in IVC1 media. After 48 hours, 50% of the media was replaced with IVC2 media. 50% of IVC2 media was refreshed daily until conclusion of the experiment.

### scRNA-seq sample preparation

Single cells were obtained upon enzymatic dissociation from day 7 and day 10 (Geltrex 5%) *in vitro* attached *blastoids*. Two different methods were used for sample preparation according to the presence of extracellular matrix in the media. For day 7 *blastoids*, treatment with Accutase (Stemcell Tecnologies) for 10 minutes at 37 degrees was sufficient to detach and isolate a single cell suspension. In order to remove the extracellular matrix from the day 10 sample, we first applied Dispase solution (Stemcell Tecnologies) for 7 minutes, washed twice with PBS^-/-^ and then applied Papain/DNase solution (Worthington) for 15 minutes at 37 degrees. Mechanical trituration followed by two washes with BSA 0.04% in PBS^-/-^ was used to isolate single cells. Before Gem formation, samples were filtered with a 40um strainer. Single Cell 3′ v.3 kit (10X genomics) was used for library preparation according to manufacturer instructions.

### scRNA-seq analysis

10X genomics libraries were sequenced using a NovaSeq 6000 (Illumina) and FASTQ files were aligned using Cell Ranger (7.0.0) against hg38 reference genome. Cell Ranger count matrices were processed in Python using Scanpy (Wolf et al., 2018) (scanpy=1.9.1, https://pypi.org/project/scanpy/). Time points of analysis were 7 dpa and 10 dpa supplemented with extracellular matrix. Counts matrices were filtered using the following criteria: minimum of 300 genes per cell and genes expressed in at least 3 cells. Cells with more than 20% of mitochondrial genes were discarded. After stringent filtering, 7354 cells (7 dpa) and 5489 cells (10 dpa) were selected for further analysis. Total-counts were normalized to 10000 reads per cell and log-transformed. The top 3000 high variable genes were used for dimensionality reduction and further analysis. Leiden clustering and differentially express genes (Wilcoxon rank-sum) were used to annotate the clusters (Table 1). Data from the two time points (7 and 10 dpa) were integrated applying scanoroma (Hie et al., 2019) (scanorama.integrate_scanpy) using commonly detected high variable genes. The *blastoids* dataset (7 and 10 dpa) or selected populations were integrated with *in vivo* human (Tyser et al., 2021; Xiang et al., 2019), mouse (Pijuan-Sala et al., 2019) and monkey (Ma et al., 2019; Zhai et al., 2022) datasets applying the scanoroma algorithm using commonly detected high variable genes. Classification using the Embryogenesis Prediction tool was performed by submitting the raw counts matrix of 5000 randomly sampled cells for each dataset (7 and 10 dpa) (Zhao et al., 2021)(https://petropoulos-lanner-labs.clintec.ki.se/app/shinyprediction). Classification results were plotted as individual colors in the UMAP space and are available in Table 2.

Raw count matrices and cell annotations are available in GEO under the accession number GSE225893.

### Immunostaining

Samples were fixed with 4% PFA in PBS^+/+^ at room temperature for 30 minutes, followed by three washes with PBS^+/+^. Samples were incubated with Antibody Solution (Normal Donkey Serum 3%, Triton-X 0.2% in PBS^-/-^) for 1h before hybridization with the primary antibodies. Primary and secondary antibodies were incubated at room temperature for 1h in Antibody Solution and followed by three washes in PBS^-/-^-TritonX 0.2%. Samples were stored in Ibidi mounting solution at 4 degrees.

List of primary antibodies:

OCT4 Santa Cruz Biotechnology (sc-5279) 1:200

BRA R&D Systems (MAB20851-100) 1:500

GATA6 R&D Systems (AF1700) 1:200

GATA3 Thermo Fisher Scientific (14-9966-82) 1:100

SOX17 R&D Systems (MAB1924) 1:200

HAND1 R&D Systems (AF3168-SP) 1:200

KRT7 Abcam (ab209600) 1:100

ISL1 DSHB (39.4D5) 1:200

MIXL1 Sigma Prestige antibody (HPA005662) 1:200

SUSD2 Miltenyi Biotec (130-106-401) 1:100

SSEA4 Miltenyi Biotec (130-122-958) 1:100

KLF17 Atlas Antibodies (HPA024629) 1:200

SOX17 R&D Systems (AF2864) 1:200

GATA4 Thermo Fisher Scientific (14-9980-82) 1:100

FOXA2 Novus Biologicals (AF2400) 1:200

### Confocal imaging

Images were acquired on laser scanning confocal microscopes (Zeiss Inverted LSM 780 or Zeiss Inverted LSM 940) with ×10, ×20 dry and ×63 oil objectives.

### Images analysis

Images were shown as Maximum Intensity Projection (MIP) of z-stacks over the z-axis using ZEN black software or Fiji_V2 (Version2.1.0/1.53c). Fiji software was used for image processing, contrast, and export. The area of positive signal for individual channels was calculated on MIP images after Ilastick pixel classification (Berg et al., 2019). In brief, individual images were processed using Ilastik, and binarized images were used for area calculation. Values were scaled to the maximum value. Plots were organized in python using numpy (1.19.5), pandas (1.1.5), matplotlib (3.2.2) and seaborn (0.11.2) libraries.

---

**Suppl. Fig 1.**
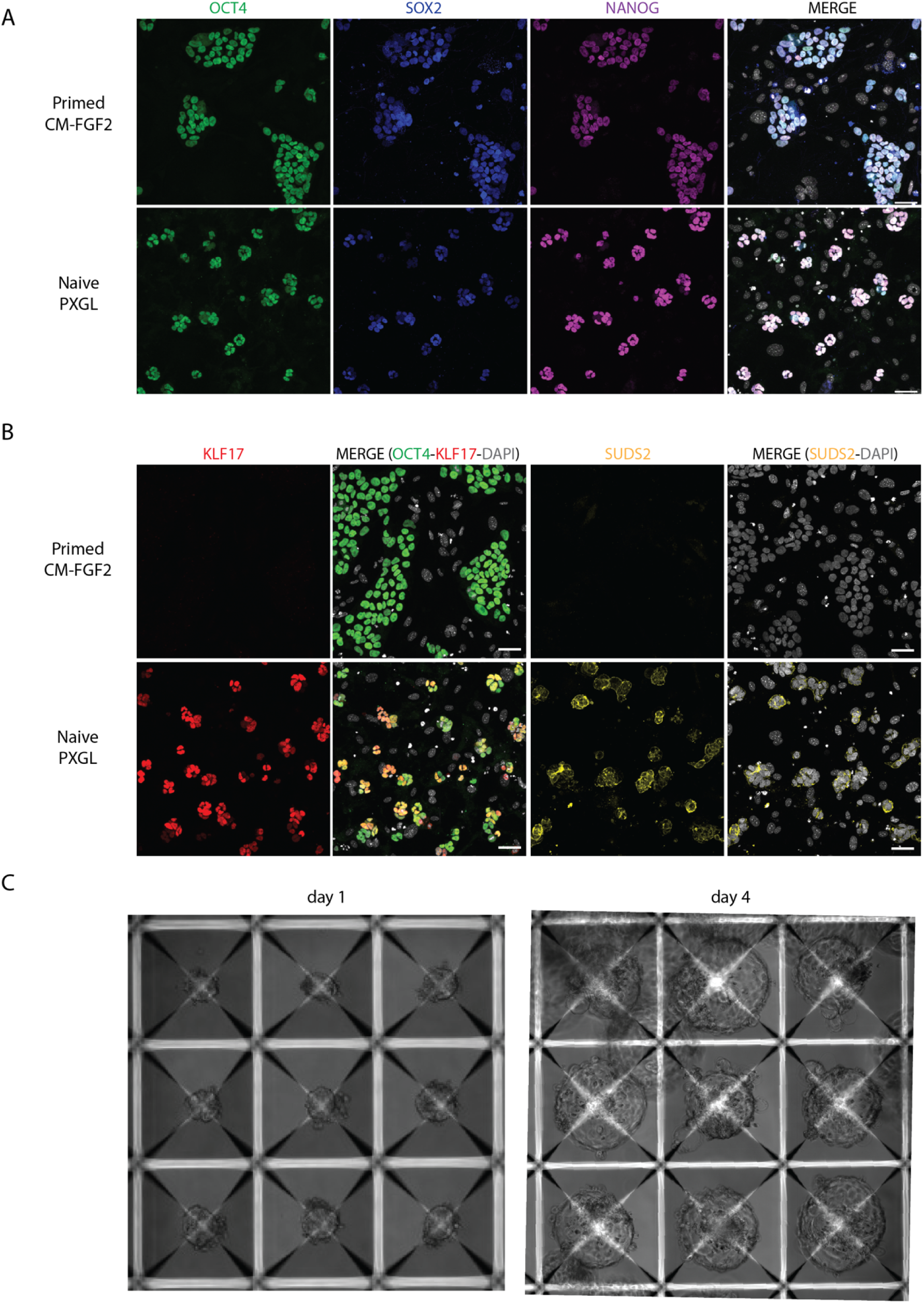
Chemically reprogrammed naïve RUES2 hESCs display preimplantation marker genes and generate *blastoids* A) Immunostaining of primed (CM+FGF2) and naïve (N2B27+PXGL) RUES2 hESCs show expression of core pluripotency genes OCT4, SOX2, and NANOG in both conditions. Immunostaining pseudo colors: OCT4 (green), SOX2 (blue) and NANOG (magenta), and DAPI (grey). Scale bar: 50um. B) Expression of the transcription factor KLF17 and surface marker SUSD2 analyzed by immunostaining distinguish naïve and primed state pluripotent stem cells. Immunostaining pseudo colors: KLF17 (red), OCT4 (green), SUDS2 (yellow), and DAPI (grey). Scale bar: 50um. C) Brightfield images during *blastoid* differentiation in Aggrewells (day 1 and day 4). Scale bar: 200um.

**Suppl. Fig 2.**
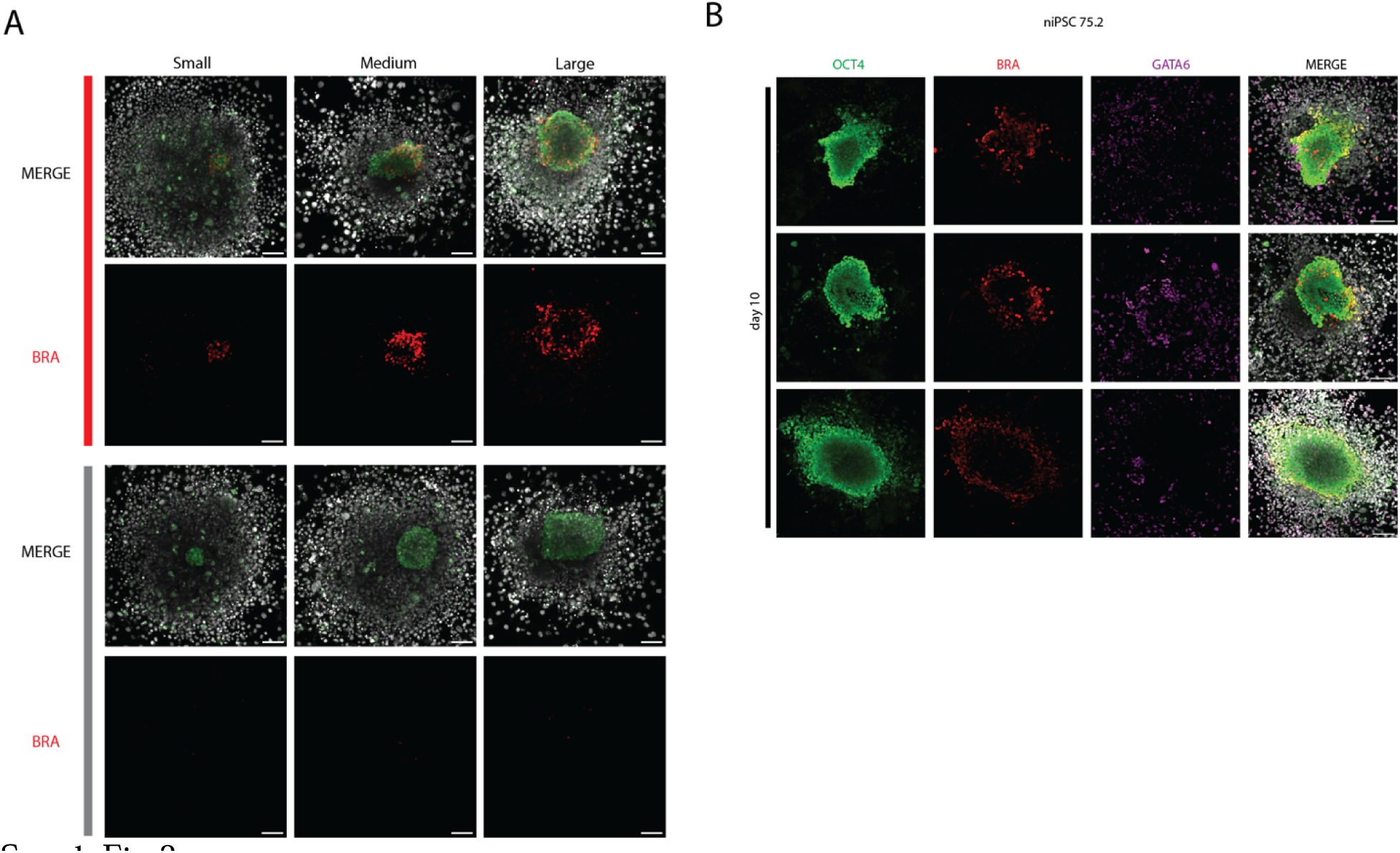
Primitive streak and mesoderm induction upon *in vitro* attachment of human *blastoids* A) Immunostaining of *in vitro* attached *blastoids* derived from at 10 dpa sorted by the size of the OCT4 positive epiblast (small, medium, and large) and whether BRA is induced (red bar = broken symmetry and grey bar = unbroken symmetry). Immunostaining pseudo colors: OCT4 (green), BRA (red), and DAPI (grey). Scale bar: 100um. B) Immunostaining of *in vitro* attached *blastoids* at 10 dpa derived from an additional pluripotent stem cell line (niPSC75.2, Bredenkamp et al., 2019). Immunostaining pseudo colors: OCT4 (green), BRA (red), GATA6 (magenta) and DAPI (grey). Scale bar: 100um.

**Suppl. Fig 3.**
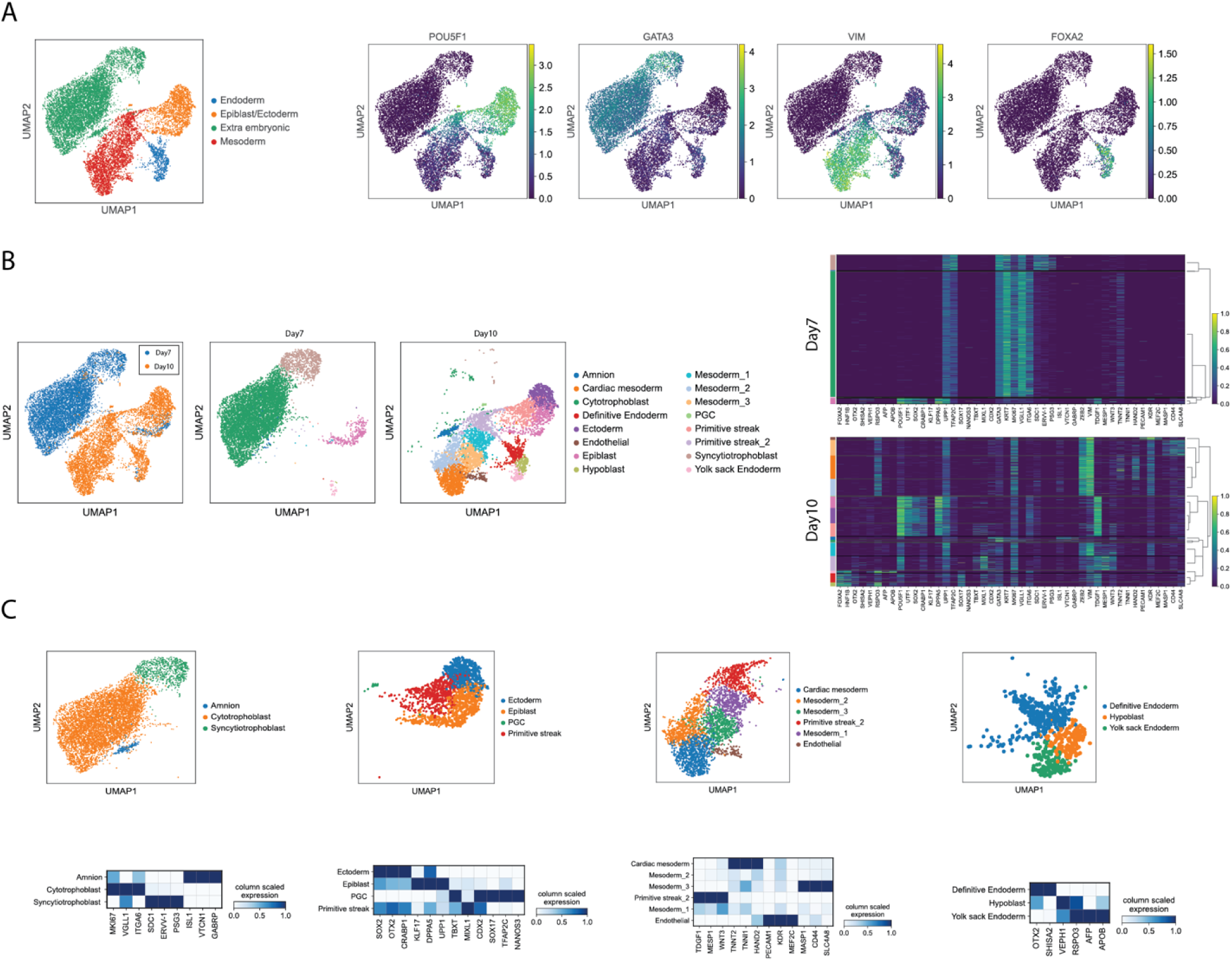
Single cell characterization of *in vitro* attached *blastoids*. A) UMAP plots display the four major cell identities and their specific marker genes: epiblast/ectoderm (POU5F1/OCT4), mesoderm (VIM), endoderm (FOXA2), and extraembryonic ectoderm (GATA3). B) UMAP plots display the identified populations at 7-10 dpa scRNA-seq dataset color coded according to the time of analysis and the cellular annotation. Heatmap of scaled counts of marker genes that distinguish individual cellular population among at 7 and 10 dpa. C) UMAP plot shows the extraembryonic sub-lineages (amnion, cytotrophoblast, and syncytiotrophoblast). z-score heatmap displays marker genes that distinguish the amnion (ISL1, VTCN1, GABRP), cytotrophoblast (MKI67, VGLL1, ITGA6), and syncytiotrophoblast (SDC1, ERVV-1, PSGC3). Epiblast/ectoderm sub-lineages (ectoderm, epiblast, PGC, and early primitive streak). z-score heatmap displays marker genes that distinguish the ectoderm (SOX2, OTX2, CRABP1), epiblast (KLF17, DPPA5, UPP1), early primitive streak (TBXT, MIXL1, TBXT), and PGC (SOX17, TFAP2C, NANOS3). Mesoderm sub-lineages (cardiac mesoderm, mesoderm 1-2-3, primitive streak 2 and endothelial). z-score heatmap displays marker genes that distinguish the primitive streak 2 (TDGF1, MESP1, WNT3), cardiac mesoderm (TNNT2, TNN1, HAND2), endothelial (PECAM1, KDR, MEF2C) and Mesoderm 3 (MASP1, CD33, SLC4A8). Endodermal sub-lineages (definitive endoderm, hypoblast and yolk sack endoderm). z-score heatmap displays marker genes that distinguish the definitive endoderm (OTX2, SHISA2), hypoblast (VEPH1, RSPO3), yolk sack endoderm (AFP, APOB).

**Suppl. Fig 4.**
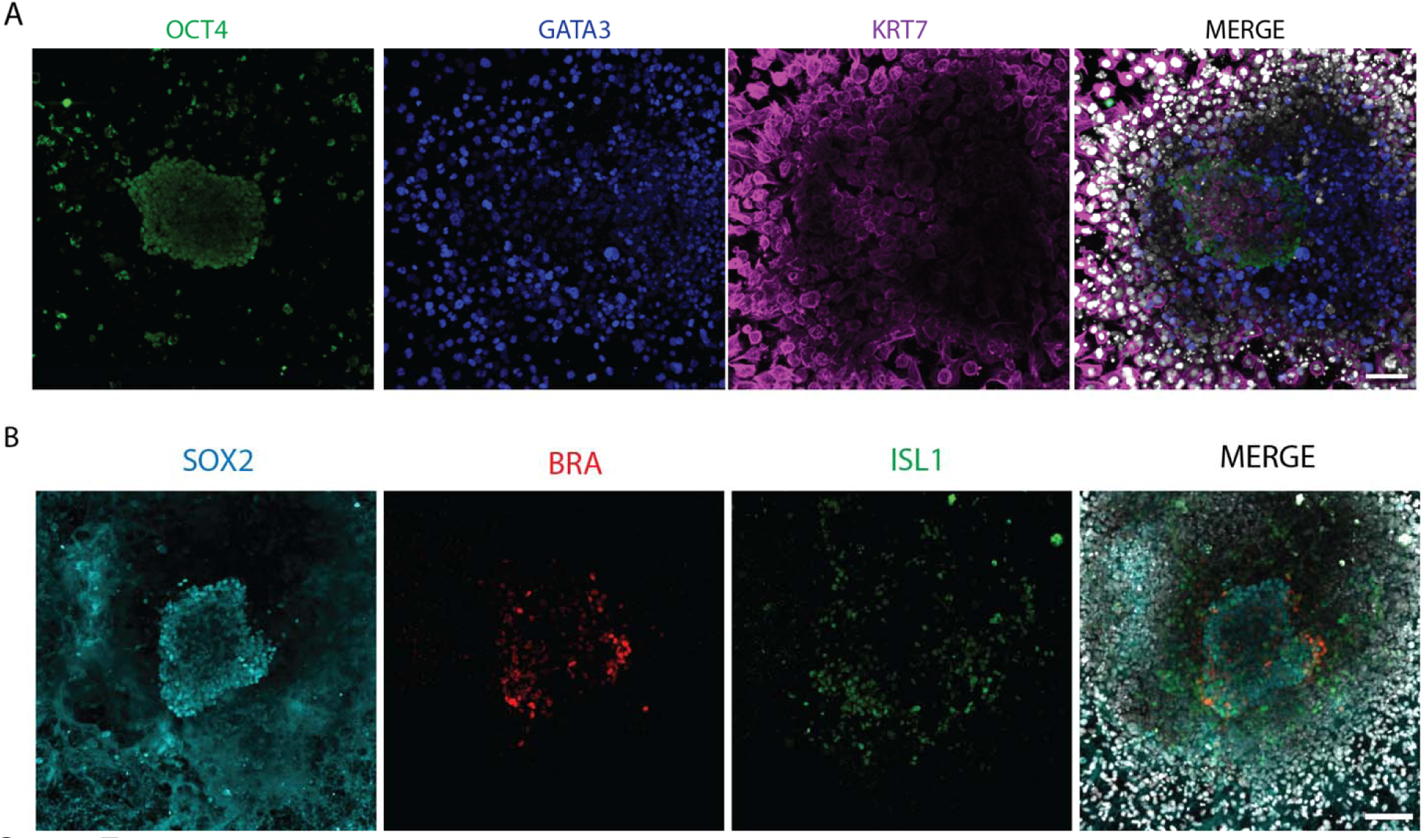
Extra-embryonic cells of the trophectoderm and the amnion at 10 dpa. A) Immunostaining of *in vitro* attached *blastoids* displayed surrounding GATA3/KRT7 positive cells at 10 dpa. The epiblast population is identified using OCT4 (green), the primitive streak/mesoderm population using BRA (red), GATA3 and KRT7 extraembryonic ectoderm population using GATA3/KRT7 (blue and magenta). Scale bar: 100um. B) Immunostaining of *in vitro* attached *blastoids* at 10 dpa. The epiblast population is identified using SOX2 (cyan), the primitive streak/mesoderm population using BRA (red), the amnion population using ISL1 (green). Scale bar: 100um.

**Suppl. Fig 5.**
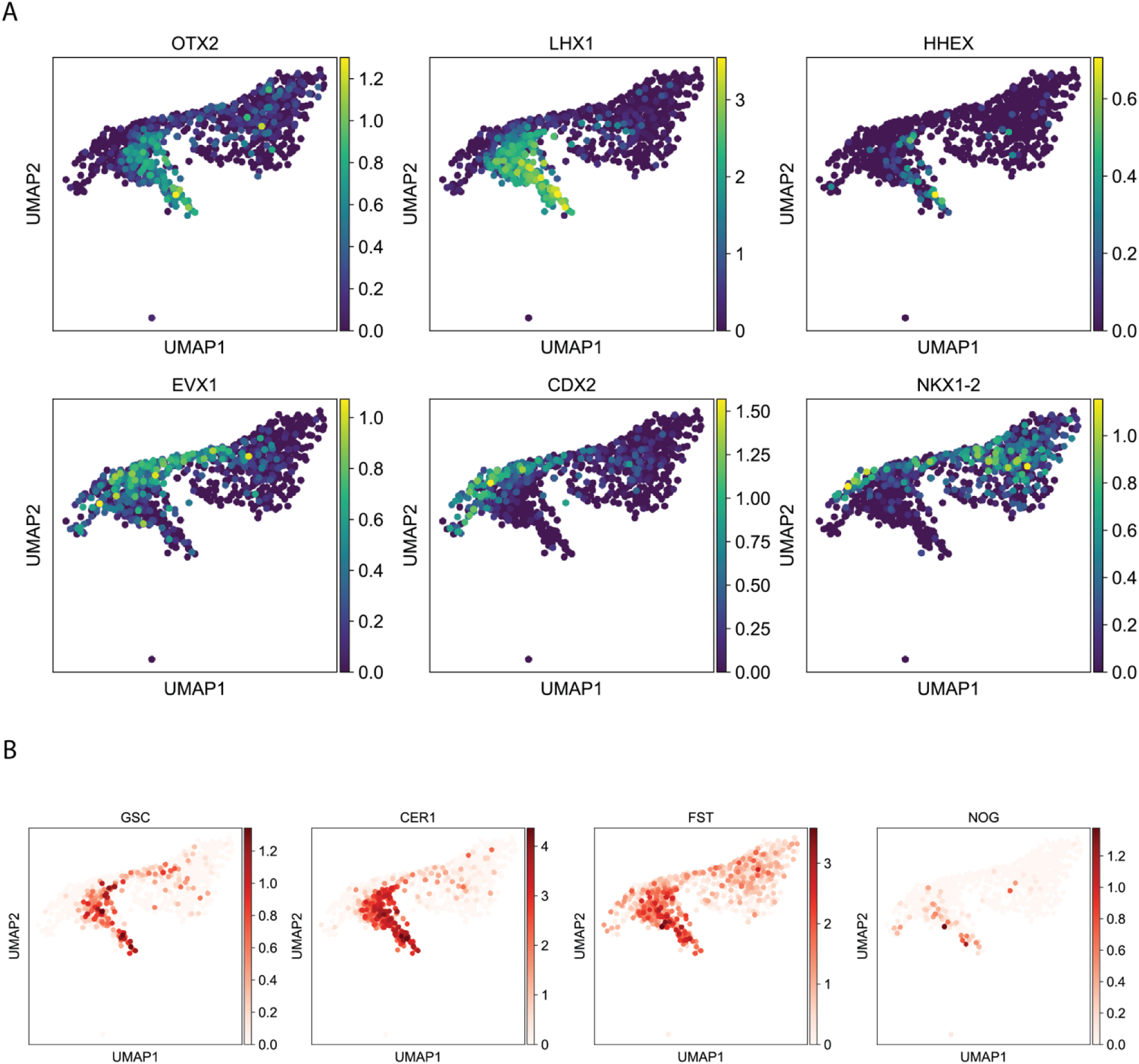
Single cell characterization of the primitive streak populations upon *in vitro* attachment A) UMAP plots of the primitive streak clusters display expression of anterior (OTX2, LHX1, HHEX) and posterior (EVX1, CDX2, NKX1-2) marker genes. B) UMAP plots show the expression of the Spemann organizer marker genes (GSC, CER1, FST, NOG).

**Suppl. Fig 6.**
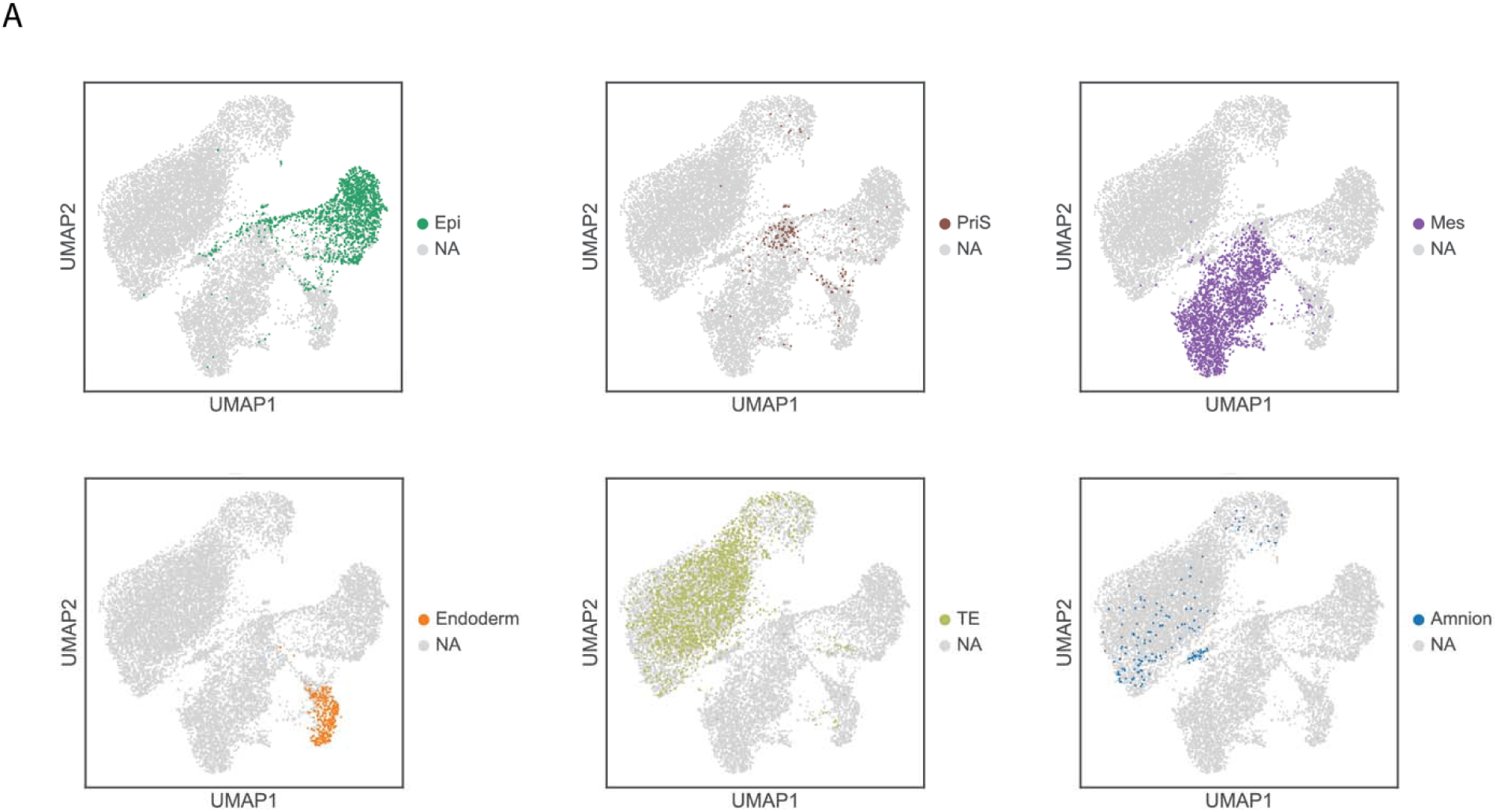
Embryonic classifier A) Predicted cell types using the Embryogenesis Prediction tool from Zhao et al., 2021 displayed as different colors in the UMAP space (color coded: epiblast, primitive streak, mesoderm, endoderm, trophectoderm, and amnion populations).

**Suppl. Fig 7:**
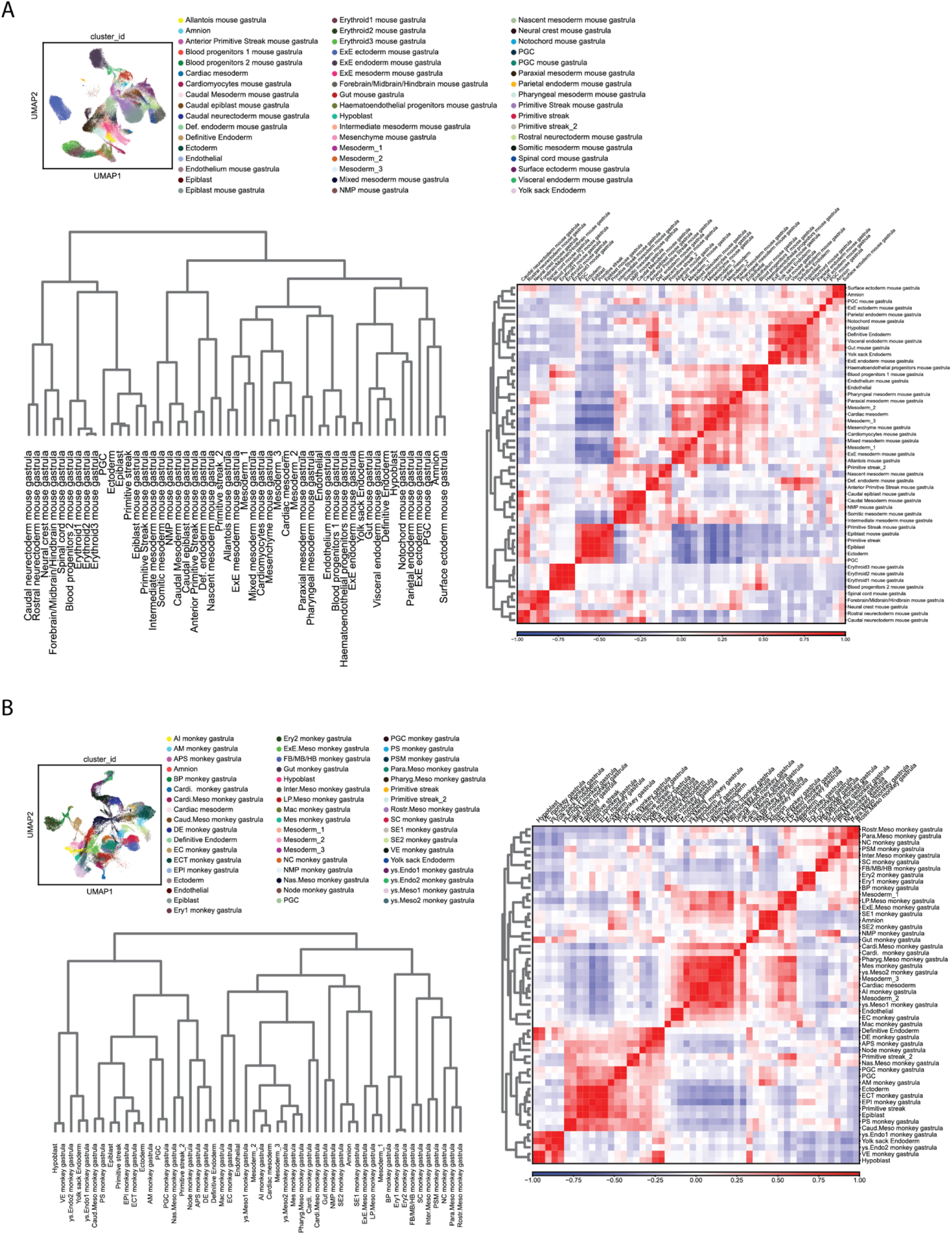
scRNA-seq datasets integration among the *in vitro* attached human *blastoids* and the *in vivo* mouse gastrulation and monkey early organogenesis datasets. D) The scRNA-seq of the in vitro attached human blastoids (7 and 10 dpa) is integrated with the mouse gastrulation dataset (Pijuan-Sala et al., 2019) using scanorma and commonly detected high variable genes. UMAP displays individual populations. The populations coming from the mouse dataset are additionally labeled as “mouse gastrula”. Correlation analysis is displayed as a z-score heatmap. Hierarchical clustering displayed as a dendrogram between the *in vitro blastoids* and *in vivo* mouse gastrula dataset. E) The scRNA-seq of the in vitro attached human blastoids (7 and 10 dpa) is integrated with the monkey early organogenesis dataset (Zhai et al., 2022) using scanorma and commonly detected high variable genes. UMAP displays individual populations. The populations coming from the mouse dataset are additionally labeled as “monkey gastrula”. Correlation analysis is displayed as a z-score heatmap. Hierarchical clustering displayed as a dendrogram between the *in vitro blastoids* and *in vivo* monkey early organogenesis dataset.

**Suppl. Fig 8:**
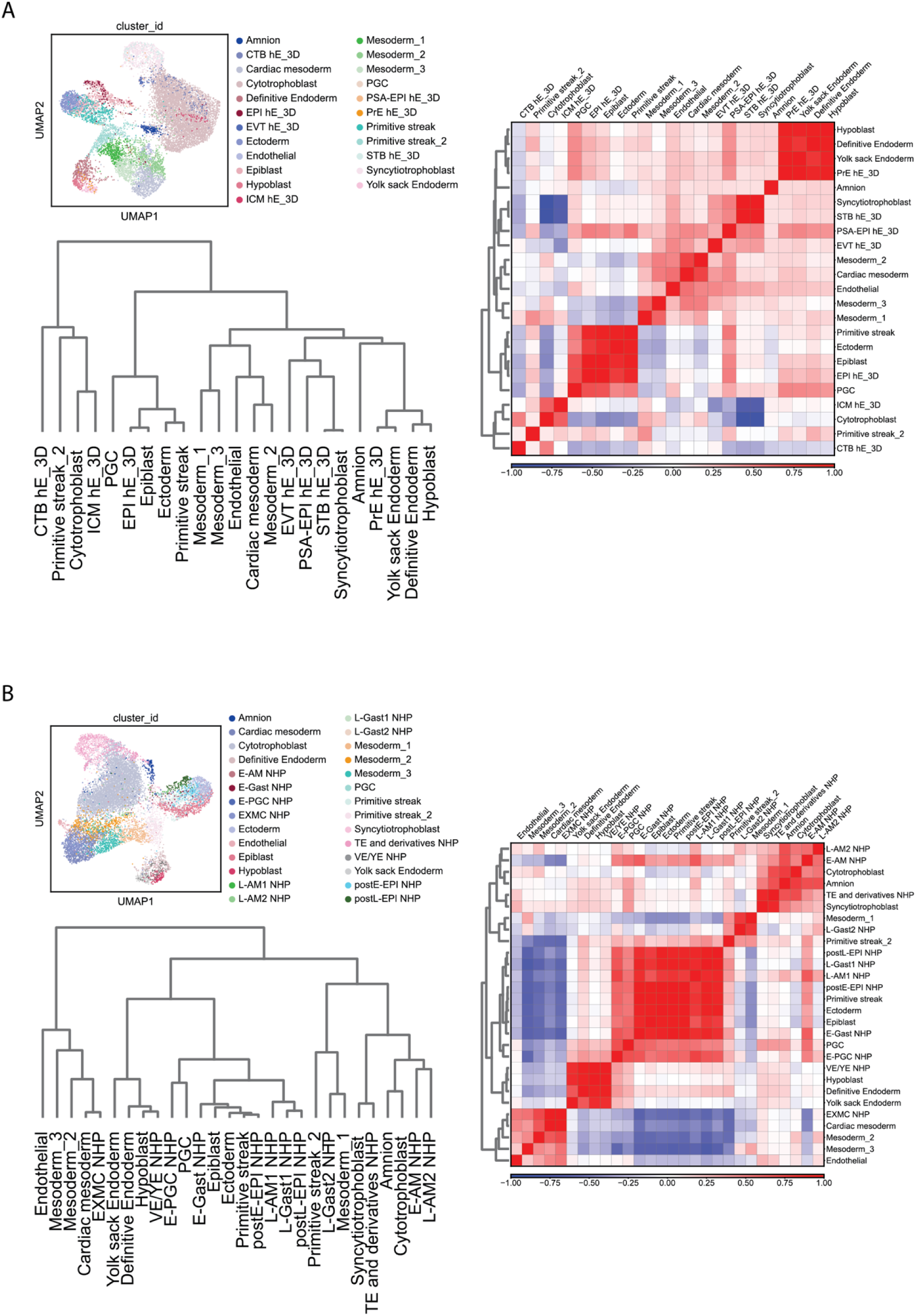
scRNA-seq datasets integration among the *in vitro* attached human *blastoids* and the *in vitro* cultured 3D human embryos and the *in vitro* attached monkey embryo datasets. A) The scRNA-seq of the in vitro attached human blastoids (7 and 10 dpa) was integrated with the human 3D cultured human embryo (Xiang et al., 2019) using scanorma and commonly detected high variable genes. UMAP displays individual populations. The populations coming from the mouse dataset are additionally labeled as “hE_3D”. Correlation analysis is displayed as a z-score heatmap. Hierarchical clustering displayed as a dendrogram between the *in vitro blastoids* and in vitro cultured human embryo in 3D. B) The scRNA-seq of the in vitro attached human blastoids (7 and 10 dpa) was integrated with in vitro attached monkey embryo dataset (Ma et al., 2019) using scanorma and commonly detected high variable genes. UMAP displays individual populations. The populations coming from the mouse dataset are additionally labeled as “NHP”. Correlation analysis is displayed as a z-score heatmap. Hierarchical clustering displayed as a dendrogram between the *in vitro blastoids* and *in vitro attached* monkey embryos.

**Suppl. Fig 9:**
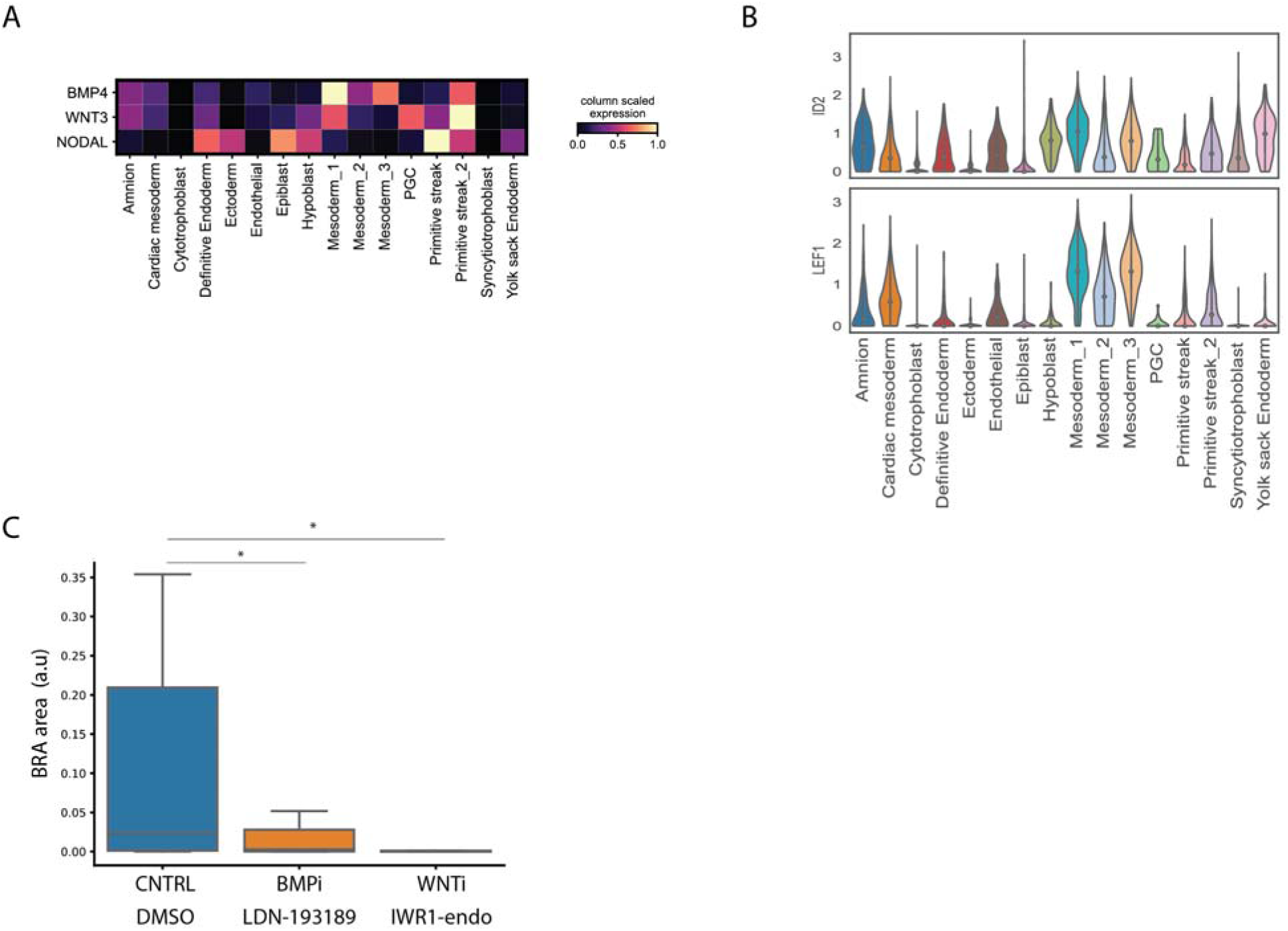
*In vitro* attached blastoids self-organize BMP, WNT and NODAL signaling pathways A) scRNA-seq analysis displayed as a heatmap of the average scaled expression of BMP4, WNT3, and NODAL ligands among individual cell populations of *in vitro* attached *blastoids*. B) scRNA-seq analysis displayed as violin plot showing the expression levels of ID2 and LEF1 which are immediate-early target genes of BMP and WNT signaling pathways, respectively. H) Immunostaining quantification of the area occupied by BRA^+^ positive cells at 10 dpa upon treatment with BMP and WNT inhibitors (LDN 193189 and IWR1-endo). Data are displayed as a boxplot (center line displays the median, and box limits are the 25th and 75th percentiles). Student’s t-test, unpaired with two tails. *p<0.05.

**Suppl. Fig 10:**
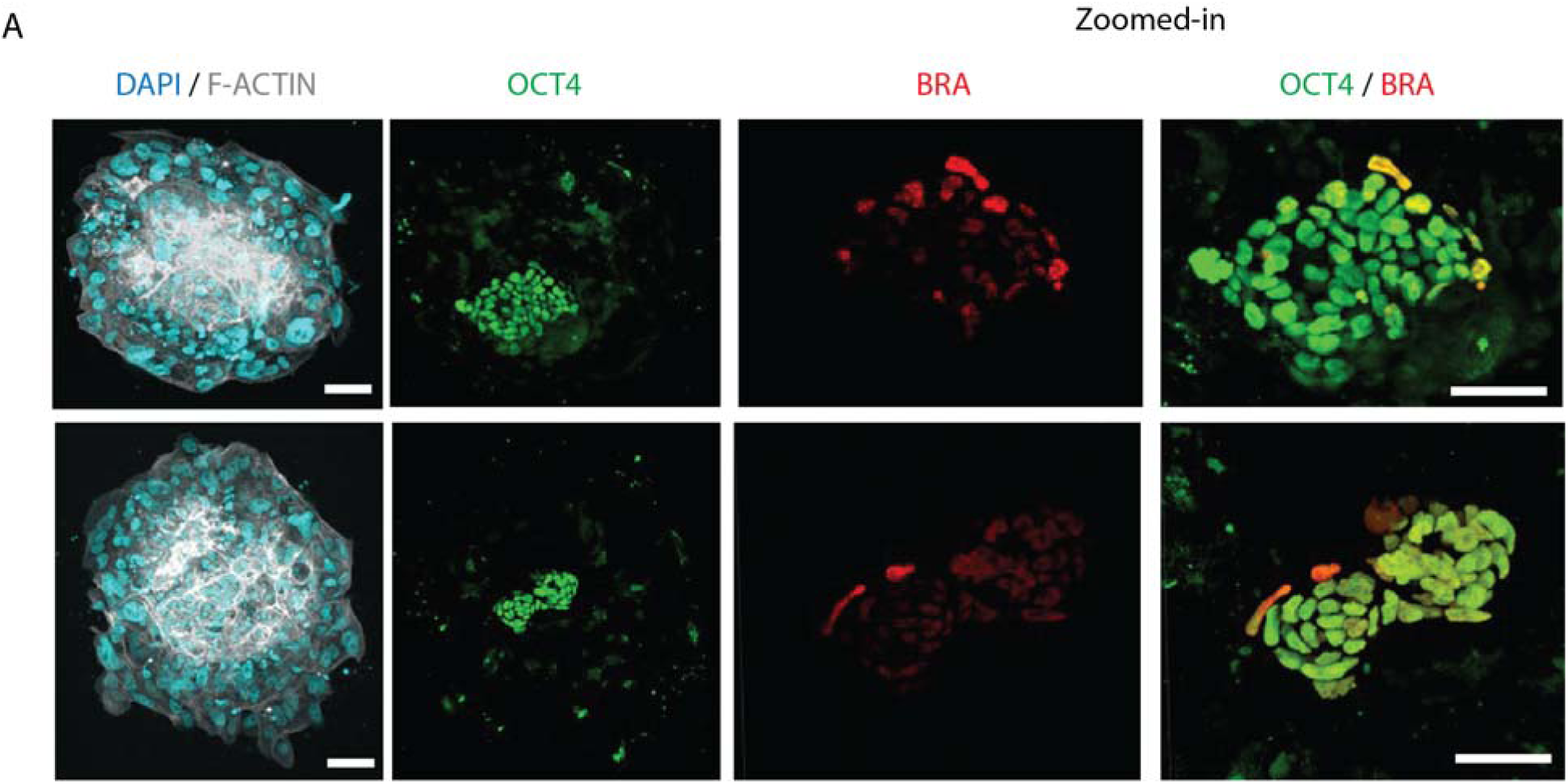
*In vitro* attached human embryos break epiblast symmetry and self-organize a BRA-positive population at 12 dpf in the absence of Laminin-521 coated plastic substrates. C) Immunostaining of *in vitro* attached human embryos at 12 dpf. Large field and zoomed-in views show DAPI (cyan), F-ACTIN (grey), OCT4 (green) and BRA (red) signals. Scale bar: 100um.

## Bibliography

1. Amadei, G., et al. Embryo model completes gastrulation to neurulation and organogenesis. Nature 610, 143–153 (2022).

2. Arnold, S. J. & Robertson, E. J. Making a commitment: cell lineage allocation and axis patterning in the early mouse embryo. Nat Rev Mol Cell Bio 10, 91–103 (2009).

3. Bredenkamp, N., Stirparo, G. G., Nichols, J., Smith, A. & Guo, G. The Cell-Surface Marker Sushi Containing Domain 2 Facilitates Establishment of Human Naive Pluripotent Stem Cells. Stem Cell Rep 12, 1212–1222 (2019).

4. Berg, S., et al. ilastik: interactive machine learning for (bio)image analysis. Nat Methods 16, 1226–1232 (2019).

5. Deglincerti, A., et al. Self-organization of the in vitro attached human embryo. Nature 533, 251– 254 (2016).

6. Gong, Y., et al. Ex utero monkey embryogenesis from blastocyst to early organogenesis. Cell 186, 2092–2110.e23 (2023).

7. Guo, G., et al. Epigenetic resetting of human pluripotency. Development 144, 2748–2763 (2017).

8. Ghimire, S., Mantziou, V., Moris, N. & Arias, A. M. Human gastrulation: The embryo and its models. Dev Biol 474, 100–108 (2021).

9. Hie, B., Bryson, B. & Berger, B. Efficient integration of heterogeneous single-cell transcriptomes using Scanorama. Nat Biotechnol 37, 685–691 (2019).

10. Irie, N., et al. SOX17 Is a Critical Specifier of Human Primordial Germ Cell Fate. Cell 160, 253– 268 (2015).

11. Jo, K., et al. Efficient differentiation of human primordial germ cells through geometric control reveals a key role for Nodal signaling. Elife 11, e72811 (2022).

12. Kagawa, H., et al. Human *blastoids* model blastocyst development and implantation. Nature 601, 600–605 (2022).

13. Ma, H., et al. In vitro culture of cynomolgus monkey embryos beyond early gastrulation. Science 366, (2019).

14. Martyn, I., Kanno, T. Y., Ruzo, A., Siggia, E. D. & Brivanlou, A. H. Self-organization of a human organizer by combined Wnt and Nodal signalling. Nature 558, 132–135 (2018).

15. Martyn, I., Brivanlou, A. H. & Siggia, E. D. A wave of WNT signaling balanced by secreted inhibitors controls primitive streak formation in micropattern colonies of human embryonic stem cells. Development 146, dev172791 (2019).

16. Minn, K. T., et al. High-resolution transcriptional and morphogenetic profiling of cells from micropatterned human ESC gastruloid cultures. Elife 9, e59445 (2020).

17. Moris, N., et al. An in vitro model of early anteroposterior organization during human development. Nature 582, 410–415 (2020).

18. Morgani, S. M., Metzger, J. J., Nichols, J., Siggia, E. D. & Hadjantonakis, A.-K. Micropattern differentiation of mouse pluripotent stem cells recapitulates embryo regionalized cell fate patterning. Elife 7, e32839 (2018).

19. Niu, Y., et al. Dissecting primate early post-implantation development using long-term in vitro embryo culture. Science 366, (2019).

20. Pijuan-Sala, B., et al. A single-cell molecular map of mouse gastrulation and early organogenesis. Nature 566, 490–495 (2019).

21. Rivera-Pérez, J. A. & Magnuson, T. Primitive streak formation in mice is preceded by localized activation of Brachyury and Wnt3. Dev Biol 288, 363–371 (2005).

22. Sasaki, K., et al. The Germ Cell Fate of Cynomolgus Monkeys Is Specified in the Nascent Amnion. Dev Cell 39, 169–185 (2016).

23. Shahbazi, M. N. Mechanisms of human embryo development: from cell fate to tissue shape and back. Development 147, dev190629 (2020).

24. Shahbazi, M. N., et al. Self-organization of the human embryo in the absence of maternal tissues. Nat Cell Biol 18, 700–708 (2016).

25. Shao, Y., et al. A pluripotent stem cell-based model for post-implantation human amniotic sac development. Nat Commun 8, 208 (2017).

26. Simunovic, M., et al. A 3D model of a human epiblast reveals BMP4-driven symmetry breaking. Nat Cell Biol 21, 900–910 (2019).

27. Simunovic, M., Siggia, E. D. & Brivanlou, A. H. In vitro attachment and symmetry breaking of a human embryo model assembled from primed embryonic stem cells. Cell Stem Cell 29, 962–972.e4 (2022).

28. Solnica-Krezel, L. & Sepich, D. S. Gastrulation: Making and Shaping Germ Layers. Annu Rev Cell Dev Bi 28, 687–717 (2012).

29. Tarazi, S., et al. Post-gastrulation synthetic embryos generated ex utero from mouse naive ESCs. Cell 185, 3290–3306.e25 (2022).

30. Tyser, R. C. V., et al. Single-cell transcriptomic characterization of a gastrulating human embryo. Nature 600, 285–289 (2021).

31. Warmflash, A., Sorre, B., Etoc, F., Siggia, E. D. & Brivanlou, A. H. A method to recapitulate early embryonic spatial patterning in human embryonic stem cells. Nat Methods 11, 847–854 (2014).

32. Wolf, F. A., Angerer, P. & Theis, F. J. SCANPY: large-scale single-cell gene expression data analysis. Genome Biol 19, 15 (2018).

33. Xiang, L., et al. A developmental landscape of 3D-cultured human pre-gastrulation embryos. Nature 577, 537–542 (2020).

34. Yanagida, A., et al. Naive stem cell blastocyst model captures human embryo lineage segregation. Cell Stem Cell (2021) doi:10.1016/j.stem.2021.04.031.

35. Yang, R., et al. Amnion signals are essential for mesoderm formation in primates. Nat Commun 12, 5126 (2021).

36. Yu, L., et al. Blastocyst-like structures generated from human pluripotent stem cells. Nature 591, 620–626 (2021).

37. van den Brink, S. C. & van Oudenaarden, A. 3D gastruloids: a novel frontier in stem cell-based in vitro modeling of mammalian gastrulation. Trends Cell Biol 31, 747–759 (2021).

38. Zhao, C. et al. Reprogrammed i*Blastoids* contain amnion-like cells but not trophectoderm. doi:10.1101/2021.05.07.442980.

39. Zhai, J., et al. Primate gastrulation and early organogenesis at single-cell resolution. Nature 612, 732–738 (2022).

40. Zhai, J., et al. Neurulation of the cynomolgus monkey embryo achieved from 3D blastocyst culture. Cell 186, 2078–2091.e18 (2023).

41. Zheng, Y., et al. Controlled modelling of human epiblast and amnion development using stem cells. Nature 573, 421–425 (2019).

